# Structural analysis and construction of thermostable antifungal chitinase

**DOI:** 10.1101/2021.09.17.460877

**Authors:** Dan Kozome, Keiko Uechi, Toki Taira, Harumi Fukada, Tomomi Kubota, Kazuhiko Ishikawa

## Abstract

Chitin is a biopolymer of *N*-acetyl-D-glucosamine with β-1,4-bond and is the main component of arthropod exoskeletons and the cell walls of many fungi. Chitinase (EC 3.2.1.14) is an enzyme that hydrolyzes the β-1,4-bond in chitin and degrades chitin into oligomers. It has been found in a wide range of organisms. Chitinase from Gazyumaru (*Ficus microcarpa*) latex exhibits antifungal activity by degrading chitin in the cell wall of fungi and is expected to be used in medical and agricultural fields. However, the enzyme’s thermostability is an important factor; chitinase is not thermostable enough to maintain its activity under the actual applicable conditions. We solved the crystal structure of chitinase to explore the target sites to improve its thermostability. Based on the crystal structure and sequence alignment among other chitinases, we rationally introduced proline residues, a disulfide bond, and salt bridge in the chitinase using protein engineering methods. As a result, we successfully constructed the thermostable mutant chitinases rationally with high antifungal and specific activities. The results provide a useful strategy to enhance the thermostability of this enzyme family.

**IMPORTANCE:** We solved the crystal structure of the chitinase from Gazyumaru (*Ficus microcarpa*) latex exhibiting antifungal activity. Furthermore, we demonstrated that the thermostable mutant enzyme with a melting temperature (Tm) 6.9 °C higher than wild type (WT) and a half-life at 60°C that is 15 times longer than WT was constructed through 10 amino acid substitutions, including five proline residues substitutions, making disulfide bonding, and building a salt bridge network in the enzyme. These mutations do not affect its high antifungal activity and chitinase activity, and the principle for the construction of the thermostable chitinase was well explained by its crystal structure. Our results provide a useful strategy to enhance the thermostability of this enzyme family and to use the thermostable mutant as a seed for antifungal agents for practical use.

## INTRODUCTION

It has been reported that nearly 80% of plant pathogens are fungi (1), and their damage to crops has become a significant problem in the agricultural industry. Up to now, chemical fungicides have been extensively adopted in combating current plant diseases. However, since fungi are eukaryotes like mammals, many chemical fungicides are highly toxic to human and fungi, and their utilization has been associated with risk. Therefore, the development of fungus-specific antifungal agents is in high demand. In a pathogenic attack, plants produce pathogenesis-related (PR) proteins as a part of systemic acquired resistance (2). Plant chitinase is considered to be one of these PR proteins. Chitinase, an enzyme that degrades chitin into oligomers, has been found to exist in a wide range of organisms. Since chitin is absent in animal cells compared to fungal cells, chitinases exhibiting the antifungal activity through the enzymatic hydrolysis of fungal cell walls (3) are expected to be used as a fungus-specific antifungal agent. According to the CAZy database (http://www.cazy.org/), chitinases (EC 3.2.1.14) are divided into two families, glycoside hydrolase (GH) family 18 (GH18) and 19 (GH19) based on amino acid sequences of their catalytic regions (4). Furthermore, according to an independent classification system for plant chitinases, they are grouped into at least five classes (classes I, II, III, IV, and V) based on their domain organization and loop deletions (5).

There are many reports of antifungal activity in plant GH19 chitinases; however, reports of antifungal activity in plant GH18 chitinases are minimal. Taira et al. have screened various tropical plants that produce latex for chitinase activity (6). Among them, latex from gazyumaru (*Ficus microcarpa*), a woody flowering plant distributed in subtropical/tropical regions in Asia, showed the highest catalytic activity of all samples assayed. Moreover, gazyumaru latex exhibits strong antifungal activity. Three types of chitinase were purified from gazyumaru latex; Gazyumaru latex chitinase-A, B, and -C (GlxChiA, B, and C) belong to class III, class I, and class III chitinase, respectively. Among them, GlxChiB, a basis class I chitinase (32 kDa, p*I* 9.3), exhibited the highest antifungal activity (6). GlxChiB consists of two domains, carbohydrate-binding module family 18 (CBM18) as a chitin-binding domain (ChBD) and GH19 domain as a catalytic domain (7). The optimum temperature for GlxChiB was 50 – 60 °C, but this enzyme was unstable above 60 °C (6).

For industrial use of the enzyme, its high activity and thermostability are important factors. The thermostability of many enzymes has been improved by protein engineering methods for their industrial use. Random mutagenesis is a typical tool for increasing thermostability (8-10). Recently computational design can be applied to the protein design (11, 12). Compared with such trial-and-error methods, rational protein design using the protein model or structural data is developing as a powerful and efficient tool for constructing thermostable enzymes (13-15). In this study, we solved the crystal structure of the catalytic domain of both WT and a thermostable mutant of GlxChiB, which was rationally designed by the protein engineering method not to affect the enzyme activity. Furthermore, based on the crystal structures of the WT and the thermostabilized enzyme, the mechanism of each mutation and new insight on the thermostabilizing effect are discussed.

## RESULTS

### Structural aspect and crystal structure of the chitinase

GlxChiB was found to belong to the GH19 family containing the chitin-binding domain (ChBD) at the N-terminus and exhibited high catalytic activity on chitin degradation and antifungal activities (7). Multiple sequence alignments of GH19 chitinases from the plant are shown in Figure 1. Only the truncated catalytic domain of WT GlxChiB (CD-WT; Amino acid numbers 45-289) was the object for the crystal structure determination in this study due to the low molecular weight ChBD (amino acid numbers 1-39) containing four disulfide bonds appearing to be a thermostable protein as observed in hevein, a small disulfide-rich protein from *Hevea brasiliensis* (16). The recombinant catalytic domain of the enzyme was prepared and purified as described previously (7). Crystals approximately 200 × 200 × 200 µm in size were obtained after more than one week at 298 K (see Materials and methods for the crystallization conditions). The diffraction data set was collected at a wavelength of 1.000 Å at the KEK PF BL5A. Data collection and refinement statistics are shown in Table 1. The diffraction data were collected up to 1.6 Å resolution. The crystals belonged to orthorhombic of space group P2(1)2(1)2(1), with unit cell *a* = 90.900 Å, *b* = 106.583Å, *c* = 107.656 Å. The initial phase of the structure was determined by molecular replacement with Phaser in the CCP4 package using the crystal structure of chitinase (GH19) from *Vigna unguiculata* (PDB: 4TX7) (17) as the search model. In the asymmetric unit, four molecules of CD-WT were identified; the Matthews coefficient (VM) (18) was calculated as 2.32 Å^3^ Da^−1^, with a solvent content of 47% (v/v). After refinement using Refmac5, R factors of the model were estimated as Rwork 22.7% and Rfree 25.3%. The value of Rfree was obtained from a test set consisting of 5% of all reflections. Ramachandran plot (19) for the model showed that 98% of the residues were in the most favored regions, with 2% of the residues in additional allowed regions. Trp121 is in Ramachandran outliners regions, with unambiguous electron density. The coordinates and structure factors for CD-WT were deposited in the Protein Data Bank under the accession code 7V91. The overall structures of four molecules of GlxChiB were built from 45 to 288 residues (Figure 2A left), and the root means square deviation (RMSD) of the Cα atoms was less than 0.457 Å over 300 Cα atoms among them. The RMSD value indicated that the overall structures of the four molecules of GlxChiB were almost identical.

**Table 1.** Data Collection and Refinement Statistics for CD-WT and CD-mt5.

|  | CD-WT | CD-mt5 |
| --- | --- | --- |
| PDB Entry | 7V91 | 7V92 |
| **Data collection** |  |  |
| Wavelength (Å) | 1.0000 | 1.0000 |
| Resolution (Å) | 50.00 – 1.60 (1.63-1.60) | 47.58-1.8 (1.70-1.60) |
| Space group | $P2_12_12_1$ | $C2$ |
| Unit cell dimensions (Å/°) | $a = 90.9, b = 106.58$<br>$c = 107.66$ | $a = 140.58, b = 38.89$<br>$c = 177.74, \beta = 96.03$ |
| No. of total reflections | 1,000,279 | 799,385 |
| No. of unique reflections | 137,756 | 235,922 |
| Rmerge*(%) | 6.1 (38.1) | 5.0 (39.9) |
| Completeness (%) | 99.8 (99.7) | 96.6 (88.5) |
| Redundancy | 7.3 (3.8) | 3.4 (3.16) |
| Average I/σ (I) | 46.0 (5.6) | 12.0 (2.3) |
| molecules/ASU | 4 | 4 |
| Matthews coefficient (Å<sup>3</sup>/Da) | 2.5 | 2.32 |
| Solvent (%) | 51 | 47 |
| **Refinement** |  |  |
| Rwork**/Rfree*** (%) | 22.7/25.3 | 21.3/24.3 |
| RMS deviations from ideal values |  |  |
| Bond angle (°) | 1.400 | 1.365 |
| Bond length (Å) | 0.006 | 0.006 |
| Ramachandran plot |  |  |
| Favored (%) | 98 | 97 |
| Allowed (%) | 2 | 2 |
| Outliers (%) | 0 (4 residues) | 0 (2 residues) |

The numbers in parentheses are for the outer shells.
\* $R_{\text{merge}} = \sum_{hkl} \sum_j |I_{j(hkl)} - \langle I_{(hkl)} \rangle| / \sum_{hkl} \sum_j I_{j(hkl)}$ , where $I_{j(hkl)}$ is the $j$ th measurement of reflection $hkl$ , including symmetry-related observations of a unique reflection, and $\langle I_{(hkl)} \rangle$ is the average.
\*\* $R_{\text{work}} = \sum |F_o - F_c| / \sum F_o$ calculated for the working set.
\*\*\* $R_{\text{free}} = \sum |F_o - F_c| / \sum F_o$ calculated for the test set (approximately 5% of reflections that were excluded from the refinement).

**Figure 1.**
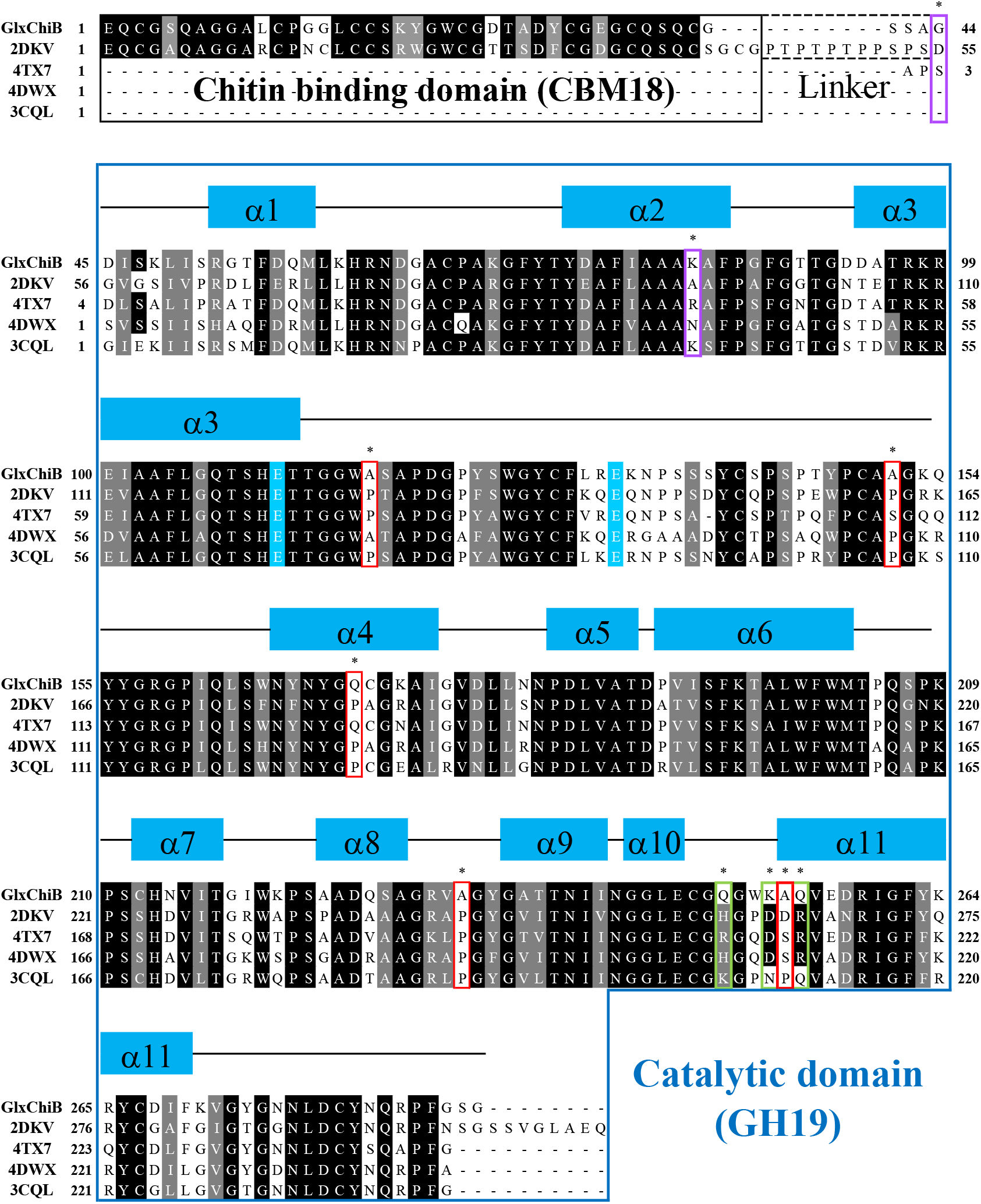
A multiple sequence alignment of GH19 chitinases from plant. Strictly conserved and similar residues are shown in white with a black and gray background, respectively. Two glutamic acid residues, which are conserved among GH19 chitinases and constructed catalytic sites are indicated with a blue background. α-helices and loops are represented as blue squares and black lines, respectively. All five proline mutation sites, two cysteine mutation sites, and mutation sites for salt bridges are indicated with red boxes, purple boxes, and green boxes, respectively. All target sites are indicated with *. The abbreviations of proteins are used for this alignment as PDB ID as follows: 2DKV, class-I chitinase from *Oryza sativa*; 4TX7, catalytic domain of class I chitinase from *Vigna unguiculata*; 4DWX, class II chitinase from *Secale cereal* (rye) seed; 3CQL, class II chitinase from *Carica papaya* (Papaya).

**Figure 2.**
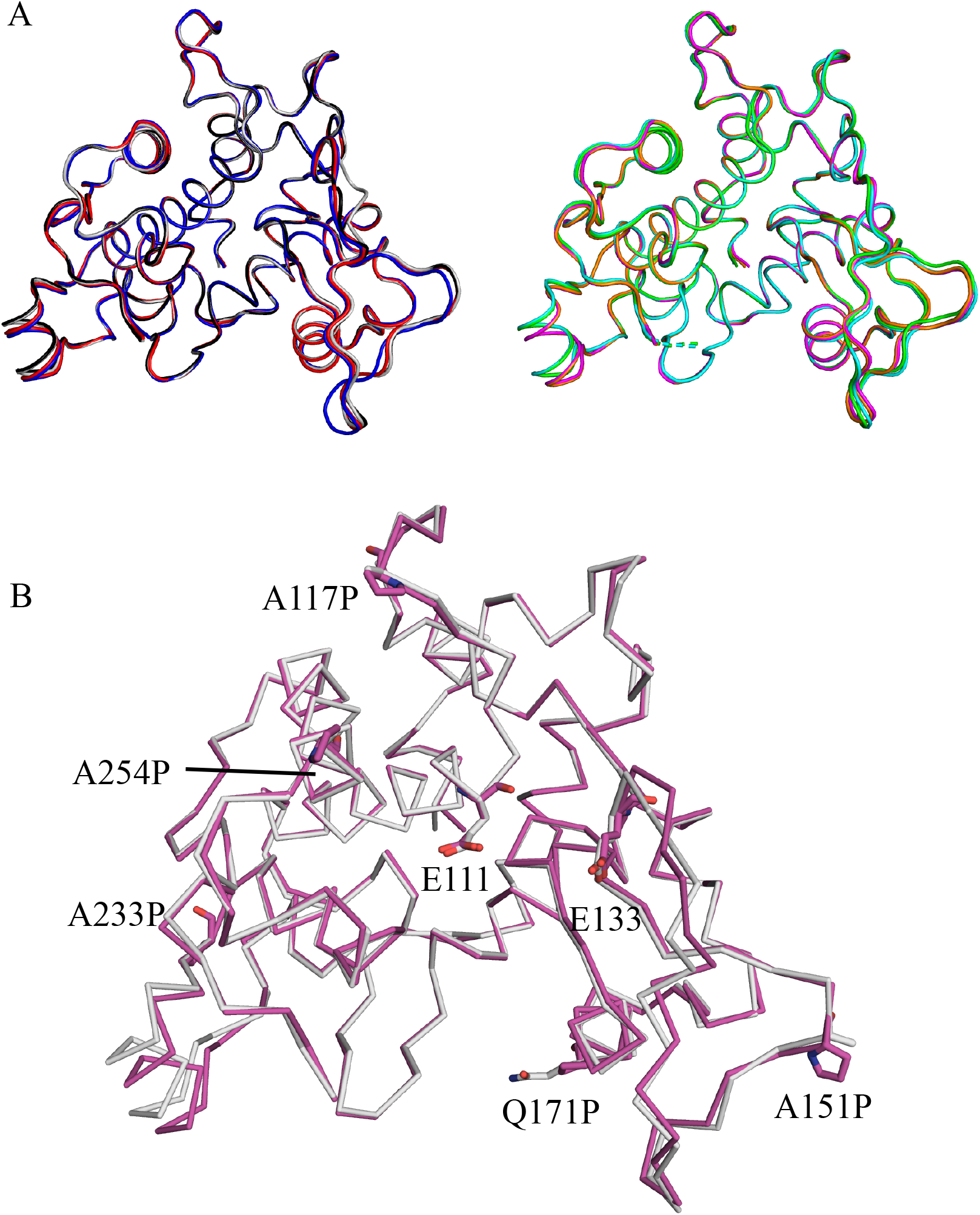
Crystal structure of GlxChiB CD-WT and CD-mt5. (A) Superimposed models of four molecules of CD-WT (left) and CD-mt5 (right). Chains A-D of CD-WT and CD-mt5 are shown as cartoon loop models and colored gray, red, blue, black, magenta, cyan, orange, and green, respectively. (B) Structural alignment of CD-WT (gray) and CD-mt5 (magenta). Cα of them are shown as ribbon models. Two catalytic glutamate residues (E111 and E133) and proline substitution sites (A117P, A151P, A171P, A233P, and A254P) are shown in stick models with atomic elements colors (O atoms, red; N atoms, blue; C atoms, gray in CD-WT and magenta in CD-mt5, respectively).

### Design of thermostable mutants of GlxChiB

Sequence alignment among the chitinases; class-I chitinase from *Oryza sativa* (PDB ID: 2DKV) (20), catalytic domain of class I chitinase from *Vigna unguiculata* (PDB ID:4TX7) (17), class II chitinase from *Secale cereale* (rye) seed (PDB ID:4DWX) (21), and class II chitinase from *Carica papaya* (PDB ID:3CQL) (22) are shown in Figure 1. In the catalytic domains of chitinase sequences, catalytic residues, disulfide bonds, and overall structures are well conserved. Proline is the only amino acid with a secondary amine, in that the side chain is directly connected to nitrogen of the main chain, preventing the rotation of phi angles of the peptide bond. Therefore, proline residues in proteins make the main chain at loop regions rigid, and several of them should be critical to the stability, including the thermostability of the protein. Furthermore, it has been proposed that the thermostability of a protein can be increased by introducing a proline that decreases the configurational entropy of unfolding (23). In the catalytic domains of GlxChiB sequence, some residues where proline residue is conserved in some other chitinases are replaced with other amino acids (Figure 1). Therefore, the introduction of proline residues in CD-WT appears tolerant and responsible for increasing its thermostability. We first took notice of the proline residues for the construction of thermostable mutants. The five positions (Ala117, Ala151, Gln171, Ala233, and Ala254) in which proline residue are not conserved in CD-WT among these enzymes (Figure 1 and 3) are remarkable points for the mutations. Referring to the crystal structure of CD-WT, Ala117 is located at the second site of a beta-turn I (Figure 4A), Ala151 is located at the second site of a beta-turn II (Figure 4B), Gln171 is located in in the middle of the helix structure (Figure 4C), and Ala233 and Ala254 are located at the loop region preceding conserved helix structure (Figure 4D and E). The following five substitutive mutations were employed in CD-WT: A117P, A151P, Q171P, A233P, and A254P.

**Figure 3.**
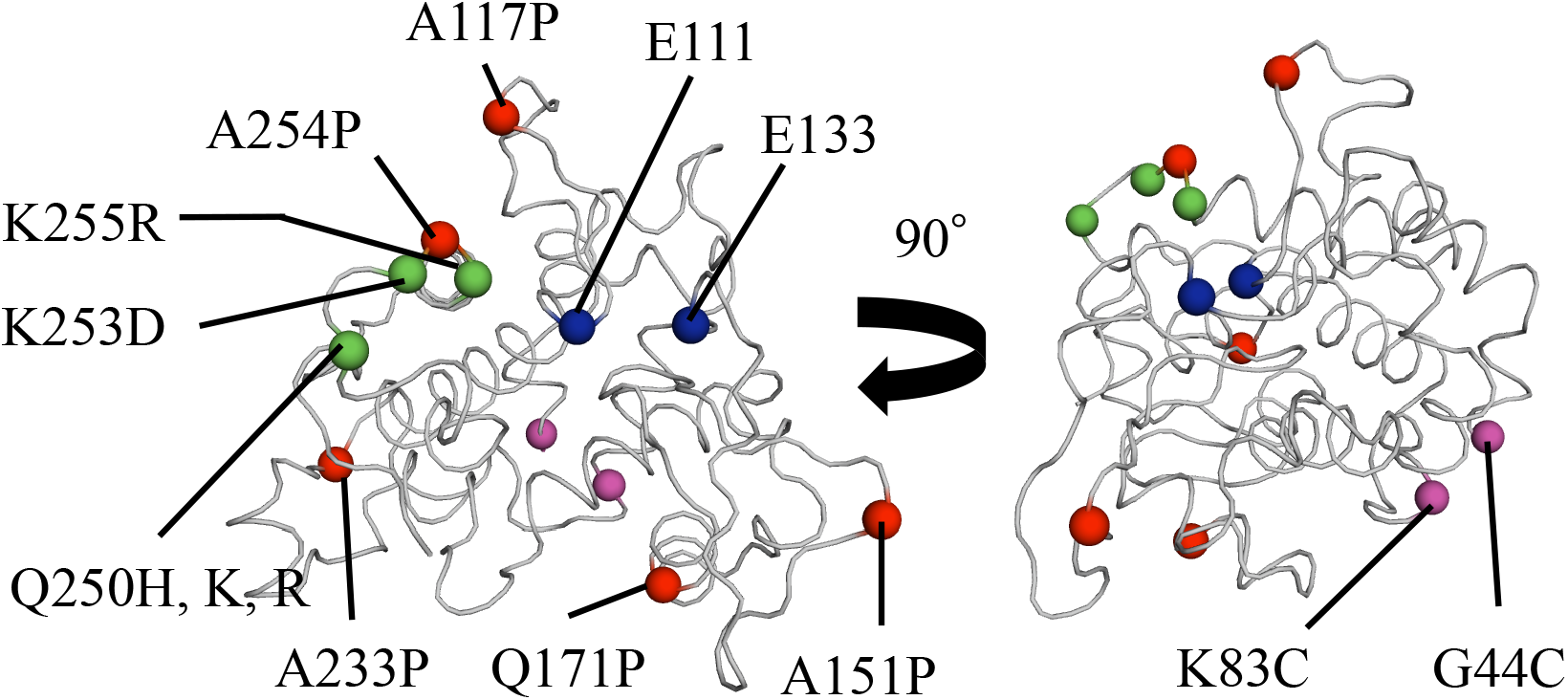
Positions of five proline substitutions and of introducing a disulfide bond and salt bridges. All five proline mutations (red sphere), two cysteine mutations (purple sphere), and three mutations for salt bridges (green sphere) are located apart from the active sites (blue sphere). Positions are labeled with the corresponding residue in WT-GlxChiB.

**Figure 4.**
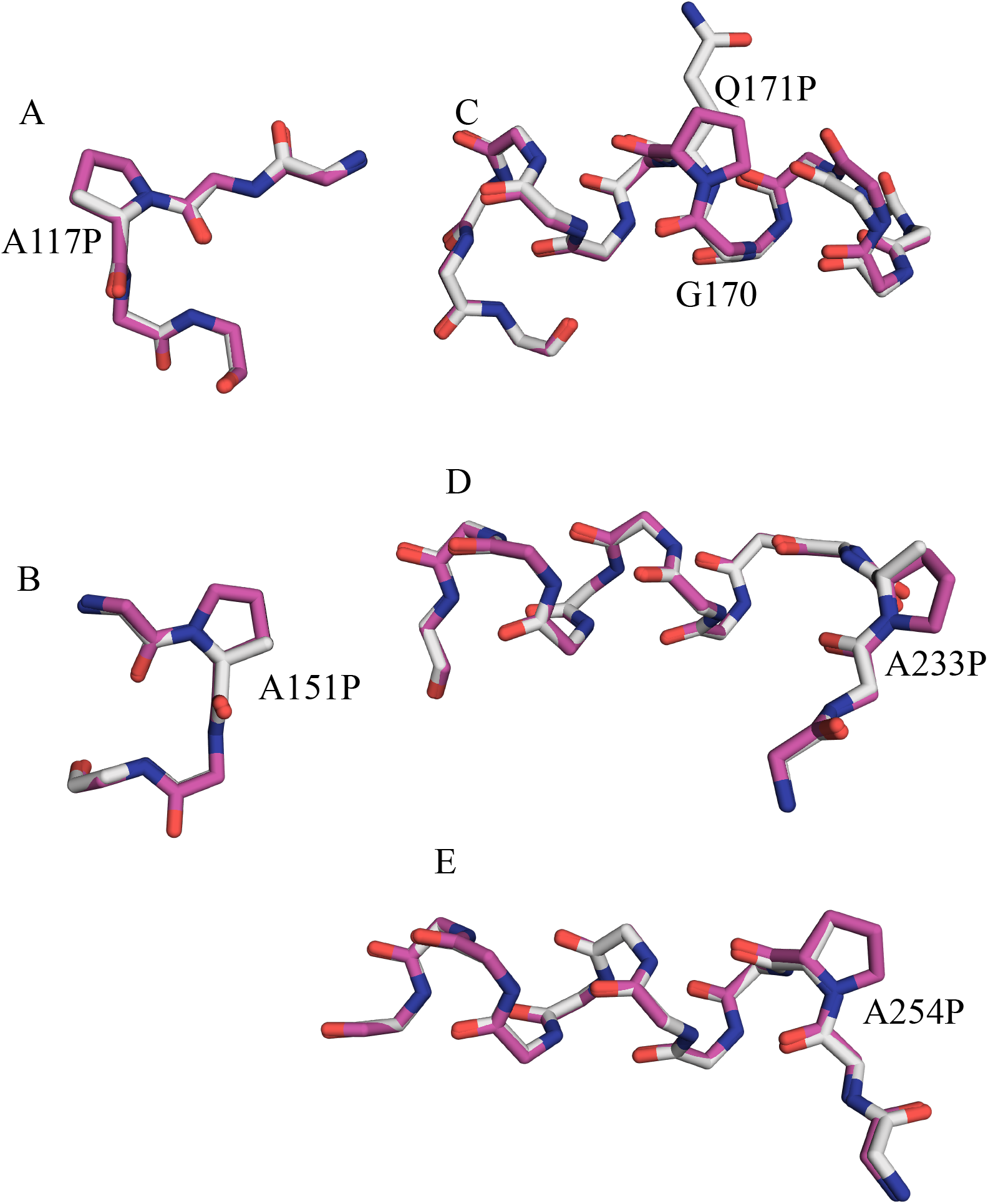
Location of mutation sites from information of crystal structure. Main chains of target regions and side chains of mutation sites are shown as a stick model with atomic elements colors (O atoms, red; N atoms, blue; C atoms, gray in CD-WT and magenta in CD-mt5, respectively). A, Ala/Pro117 is located at second site of a beta-turn I. B, Ala/Pro151 is located at second site of a beta-turn II. C, Gln/Pro171 is located in conserved helix structure. D, Ala/Pro233 is located at the loop region preceding a conserved helix structure. E, Ala /Pro254 is located at the loop region, preceding a conserved helix structure.

A disulfide bridge linking the two cysteines and their respective main peptide chains can restrict the motion of the unfolded, random coil of protein or stabilize the folded state of the protein. One disulfide bridge can contribute 2.3–5.2 kcal/mol to the thermodynamic stability of proteins (24). Considerable evidence has demonstrated the thermostability effects of engineered disulfide bridges in protein. The crystal structural analysis clarified that CD-WT contains three disulfide bonds and free Cys residues. Using the program SSBOND (25) referring to the crystal structure of CD-WT, we cannot find the suitable position in which the additional disulfide bond can be introduced in CD-WT. We have no structural information of the linker region (amino acid numbers 40-44) between ChBD and CD-WT. However, from the crystal structure, it is estimated that the distance between Cαs of N-terminal Asp45 and Lys83 is 6-8 Å in all CD-WT molecules, and then the distance between Cαs of Gly44 (in the linker region) preceding Asp45 and Lys83 is considered to be similar. Therefore, the disulfide bond between Cys44 and Cys83 could form (Figure 3) and the mutant employed in G44C/K83C. In addition, it has been reported that salt bridges play an important role in the thermostability of many proteins. The ionizable side chains frequently form ion pairs in many protein structures. Since electrostatic attraction between opposite charges is strong *per se*, salt bridges can be regarded as an important factor stabilizing the protein structure. In addition, many salt bridges were observed at the surface of thermophilic enzymes (26-28). The loop region 250-255 located at the surface loop region between two helixes appeared flexible. We targeted the loop region 250-255 located at the surface loop region between two helixes in GlxChiB (Figure 3). Significant salt bridges were observed at the corresponding region in 2DKV, 4TX7, 4DWX, and 3CQL but not in GlxChiB according to the multiple sequence alignment (Figure 1). Therefore, we tried to modify the three amino acids Q250/K253/Q255 to introduce salt bridges at the region in CD-WT by referring to these structures. Two mutants, Q250K/K253D/Q255R and Q250R/K253D/Q255R were prepared.

### Comparison of the thermostability of the mutants

Recombinant GlxChiB and their mutants described above were prepared and purified by the same method previously reported (7). The enzymatic activity assays were performed as described in the material method. Their specific activities at 37 °C were not meaningfully affected by these mutations (Table 2). WT GlxChiB exhibited the maximum activity at the temperatures around 55 °C under the conditions employed at pH 7.0 for 15 minutes (Figure S1). The thermostability was examined by the residual activities of enzymes after the incubation at 60 °C. All proline substitution mutants showed a longer half-life than that of WT (Table 2). Furthermore, as for the melting temperature (Tm) of the mutants measured by differential scanning calorimetry (DSC), all proline substitution mutants exhibited higher Tm values than that of WT (Table 2). For G44C/K83C, Tm did not change significantly; however, its half-life at 60 °C was elongated (Table 2). For the two mutants with a salt bridge network introduced (Q250K/K253D/Q255R and Q250R/K253D/Q255R), the specific activities at 37 °C were slightly increased, and the half-life at 60 °C was elongated (Table 2).

**Table 2.**
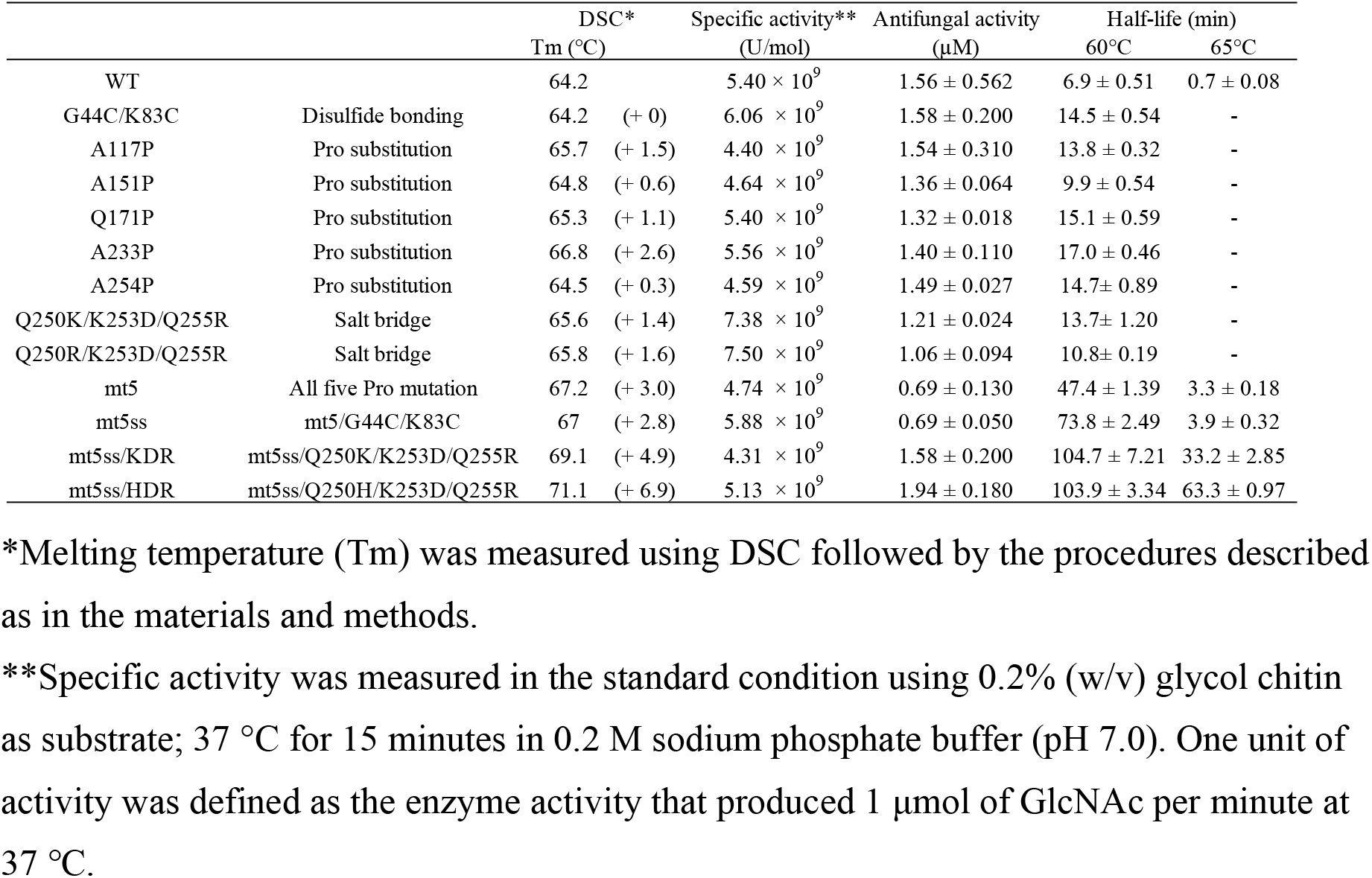
Summary of the enzymatic activities and thermodynamic parameters of the GlxChiB and its mutants.

|  |  |  | DSC* | Specific activity** | Antifungal activity | Half-life (min) |  |
| --- | --- | --- | --- | --- | --- | --- | --- |
|  |  |  | Tm (°C) | (U/mol) | (μM) | 60°C | 65°C |
| WT | | | 64.2 | $5.40 \times 10^9$ | $1.56 \pm 0.562$ | $6.9 \pm 0.51$ | $0.7 \pm 0.08$ |
| G44C/K83C | Disulfide bonding | | 64.2 (+ 0) | $6.06 \times 10^9$ | $1.58 \pm 0.200$ | $14.5 \pm 0.54$ | - |
| A117P | Pro substitution | | 65.7 (+ 1.5) | $4.40 \times 10^9$ | $1.54 \pm 0.310$ | $13.8 \pm 0.32$ | - |
| A151P | Pro substitution | | 64.8 (+ 0.6) | $4.64 \times 10^9$ | $1.36 \pm 0.064$ | $9.9 \pm 0.54$ | - |
| Q171P | Pro substitution | | 65.3 (+ 1.1) | $5.40 \times 10^9$ | $1.32 \pm 0.018$ | $15.1 \pm 0.59$ | - |
| A233P | Pro substitution | | 66.8 (+ 2.6) | $5.56 \times 10^9$ | $1.40 \pm 0.110$ | $17.0 \pm 0.46$ | - |
| A254P | Pro substitution | | 64.5 (+ 0.3) | $4.59 \times 10^9$ | $1.49 \pm 0.027$ | $14.7 \pm 0.89$ | - |
| Q250K/K253D/Q255R | Salt bridge | | 65.6 (+ 1.4) | $7.38 \times 10^9$ | $1.21 \pm 0.024$ | $13.7 \pm 1.20$ | - |
| Q250R/K253D/Q255R | Salt bridge | | 65.8 (+ 1.6) | $7.50 \times 10^9$ | $1.06 \pm 0.094$ | $10.8 \pm 0.19$ | - |
| mt5 | All five Pro mutation | | 67.2 (+ 3.0) | $4.74 \times 10^9$ | $0.69 \pm 0.130$ | $47.4 \pm 1.39$ | $3.3 \pm 0.18$ |
| mt5ss | mt5/G44C/K83C | | 67 (+ 2.8) | $5.88 \times 10^9$ | $0.69 \pm 0.050$ | $73.8 \pm 2.49$ | $3.9 \pm 0.32$ |
| mt5ss/KDR | mt5ss/Q250K/K253D/Q255R | | 69.1 (+ 4.9) | $4.31 \times 10^9$ | $1.58 \pm 0.200$ | $104.7 \pm 7.21$ | $33.2 \pm 2.85$ |
| mt5ss/HDR | mt5ss/Q250H/K253D/Q255R | | 71.1 (+ 6.9) | $5.13 \times 10^9$ | $1.94 \pm 0.180$ | $103.9 \pm 3.34$ | $63.3 \pm 0.97$ |

\*Melting temperature (Tm) was measured using DSC followed by the procedures described as in the materials and methods.
\*\*Specific activity was measured in the standard condition using 0.2% (w/v) glycol chitin as substrate; 37 °C for 15 minutes in 0.2 M sodium phosphate buffer (pH 7.0). One unit of activity was defined as the enzyme activity that produced 1 μmol of GlcNAc per minute at 37 °C.

### The integrated Pro mutant and the crystal structure of the mutant

For the thermostable proline substitute mutants (A117P, A151P, Q171P, A233P, and A254P), no decrease of the enzymatic or antifungal activity was observed (Table 2 and Figure 5). Table 2 shows that these substitutions did not affect both activities. All proline substitutions elongated half-life at 60 °C 1.4-2.4 times longer than WT, and the individual effects on thermostability were considered cumulative. To examine the structural effects of proline substitutions on the enzyme’s thermostability, we prepared the integrated mutant, mt5 (A117P/A151P/Q171P/A233P/A254P) and solved the crystal structure of the catalytic domain of mt5 (CD-mt5). CD-mt5 was prepared and purified as the same as for CD-WT. The crystals of CD-mt5 were grown under different conditions from those for CD-WT. Data collection and refinement statistics are shown in Table 1. The diffraction data was collected to a resolution of 1.80 Å. The crystals belonged to space group C2, with unit cell *a* = 140.58 Å, *b* = 38.89 Å, *c* = 177.74 Å, β=96.03°; note this the crystal system is completely different from that of CD-WT. The coordinates and structure factors for CD-mt5 were deposited in the Protein Data Bank under the accession code 7V92. The overall structures of four molecules in an asymmetric unit were built from 45 to 288 (Figure 2A, right), and the RMSD of the Cα atoms were less than 0.455 Å over 300Cα atoms among them. The RMSD value indicated that the overall structures of the four molecules of CD-mt5 were almost identical. Furthermore, the RMSD of the Cα atoms were less than 0.530 Å on average over 300Cα atoms between CD-WT and CD-mt5 in chain A. This result shows that the overall structures and the active site between WT and mt5 were almost identical. Figure 2B and Figure 4 show the structural differences between CD-WT and CD-mt5 for the five mutation sites (A117P, A151P, Q171P, A233P, and A254P) and the active site residues, Glu111 and Glu133. The structures of the main chain and side chains around the mutation points were not influenced by these mutations. This result shows that all positions for the introduction of proline residue into WT are ideal for the construction of thermostable enzymes.

**Figure 5.**
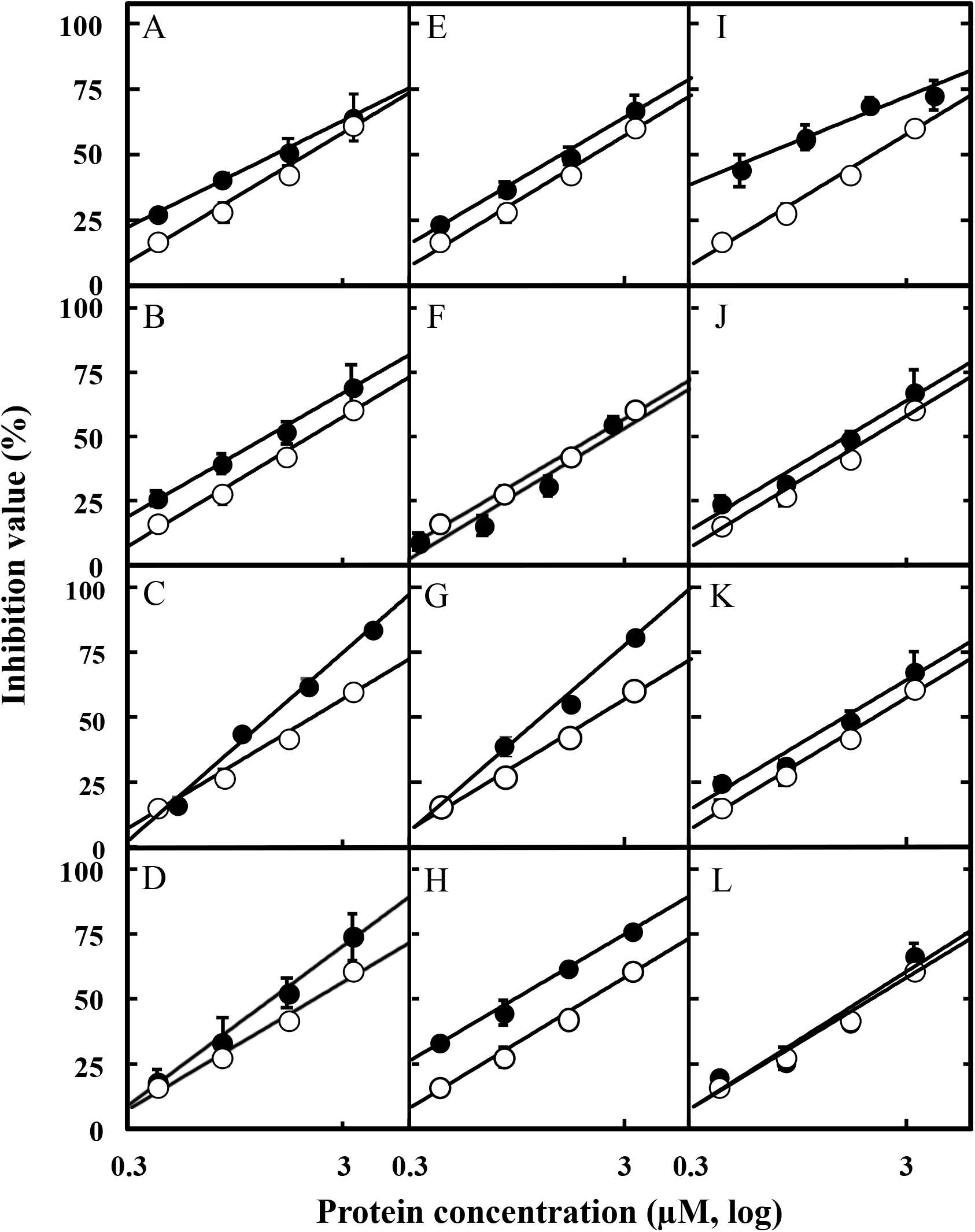
Antifungal activities of thermostable mutants. Quantitative antifungal activity assays were performed as described in the methods. Five microliters of the sample were overlaid onto an agar disk containing the mycelium of *T. viride* on a PDA plate, and then, the plate was incubated at 25 °C for 12 hours. After incubation, the re-growth area of mycelia was measured. Open and closed circles indicate wild-type and mutants, respectively. (A-E) Proline substitution: A, A117P; B, A151P; C, Q171P; D, A233P; E, A254P. (F-H) Introducing disulfide bonding and salt brides: F, G44C/K83C; G, Q250K/K253D/Q255R; H, Q250R/K253D/Q255R. (I-L) Integrated mutants: I, mt5; J, mt5ss; K, mt5ssKDR; L, mt5ssHDR. All assays were triplicate. Error bars represent ± SD (n = 3).

It was proved that the additional mutations for G44C/K83C and Q250K/K253D/Q255R into mt5 were also ideal for the construction of the thermostable enzymes by crystal structural analysis. Therefore, all positive mutations (mt5, G44C/K83C, Q250K/K253D/Q255R) were integrated and inspected. Table 2 shows that these mutations do not influence the specific activity and the individual effect for thermostability caused by mutations was additive. Half-life at 60 °C was elongated by accumulating mutations: half-life at 60 °C of mt5, mt5/G44C/K83C (mt5ss), and mt5ss/Q250K/K253D/Q255R were about 7, 11 and 15 times longer than that of WT, respectively (Table 2 and Figure 6A). mt5ss/Q250K/K253D/Q255R was proved to be the best enzyme for this study (Table 2). For the 250th position, however, histidine residue was observed in other GH19 chitinases (2DKV and 4DWX) (Figure 1). In addition, the alternative mutant mt5ss/Q250H/K253D/Q255R was examined (Table 2). There was no significant difference in half-life at 60 °C of mt5ss/Q250H/K253D/Q255R and mt5ss/Q250K/K254D/Q255R, but half-life at 65 °C of mt5ss/Q250H/K253D/Q255R was about two times longer than that of mt5ss/Q250K/K254D/Q255R (Table 2 and Figure 6B). mt5ss/Q250H/K253D/Q255R showed the highest Tm (71.1 °C: 6.9 °C higher than that of WT) among the mutants and elongated its half-life at 60 °C 15 times and at 65 °C 90 times more than that of WT, respectively (Table 2).

**Figure 6.**
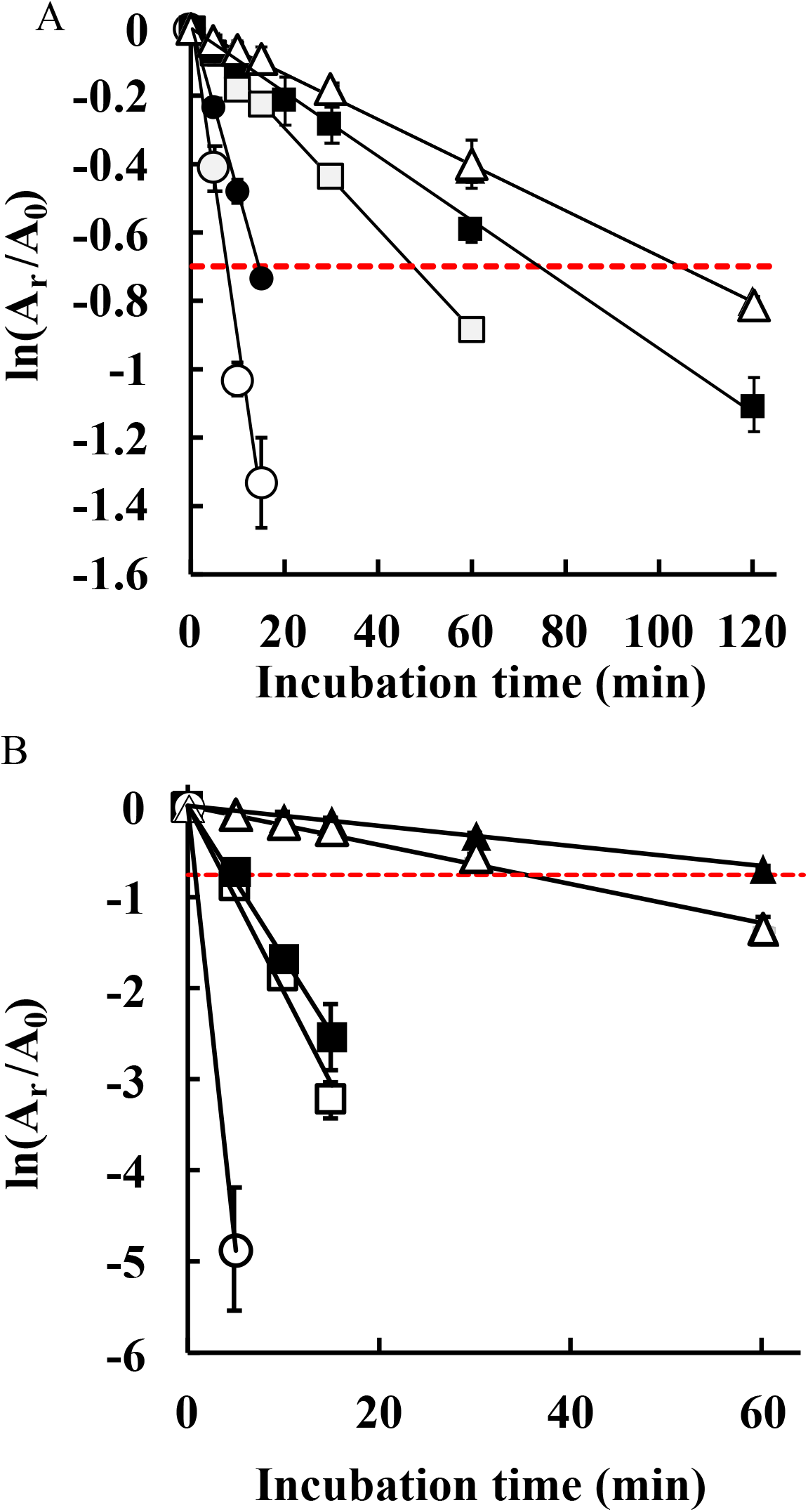
Thermostability of GlxChiB WT and mutants. Enzyme samples in a 10 mM sodium phosphate buffer (pH 7.0) were incubated at 60 °C (A) and 65 °C (B) for each period. After the treatment, the residual activities (A_r_) were measured at 37 °C for 15 minutes in a 0.2 M sodium phosphate buffer (pH 7.0), with the initial activity without heat treatment taken as 100% (A_0_). Open circle, closed circle, open square, closed square, open triangle, and closed triangle indicate WT, G44C/K83C, mt5, mt5/G44C/K83C, mt5/G44C/K83C/KDR, and mt5/G44C/K83C/HDR, respectively. Red dashed lines indicate where the value of A_r_/A_0_ = 1/2: the residual activity is 50% of initial activity. All assays were triplicate. Error bars represent ± SD (n = 3).

### Comparison of the antifungal activity of the mutants

The antifungal activities of GlxChiB and its mutants were determined by using the hyphal re-extension inhibition assay with *Trichoderma viride* as the test fungus (Figure 5). We define IC_50_ as the concentration where 50% of the hyphal re-extension areas are inhibited. As shown in Table 2 and Figure 5, the IC_50_ of WT is 1.56 ± 0.562 µM, and all mutants exhibit the same level of WT for their antifungal activities. It shows that all thermostabilized mutations do not affect their antifungal activities significantly.

## DISCUSSION

### Obtaining thermostable mutants

It is ideal for efficient antifungal activity to degrade chitin for actual use, which is realized by having the enzyme perform under various conditions. Thus, enzymes must be resistant to high temperature, organic solvents, not-neutral pH, and other chemicals. Thermostabilized chitinases are expected to exhibit excellent performance under such conditions. GlxChiB obtained from the gazyumaru latex has the highest antifungal activity among the other chitinases isolated from the latex and is expected to potentially be applied practically. While the optimum temperature for GlxChiB is 50 – 60 °C, this enzyme is unstable above 60 °C. Industrial use of this enzyme requires improving their thermostability without decreasing their enzymatic performance. However, there is often a trade-off between thermostability and enzyme activity (29-31). In this study, we successfully obtained several thermostable mutants without decreasing their activities by rational design based on sequence and structural comparison among homologous enzymes. The thermostable GlxChiB created in this study is promising and has a high potential for its application. To date, two correlations about the primary sequences of an enzyme and its thermostability have been proved. One-point mutation can improve the thermostability of an enzyme, but this effect is in general minuscule (23, 32). The second is that such effects caused by individual structural changes from one-point mutations will be cumulatively counted and added (13-15, 33). These facts suggest that an enzyme is heat adaptable (34). Several studies (35-37) provided this trend about heat adaptation; however, a fundamental principle for the design of heat adaptable mutant enzymes has not been discovered (38).

### Structural interpretation of the thermostability by Proline substitution

Proline is a unique amino acid residue in that the side chain is covalently bound to the preceding peptide bonded nitrogen and the five-membered ring imposes rigid constraints on the N-Cα rotation (39). Thus, it is proposed that the substitution of proline for appropriate amino acids can increase the stability of a protein by decreasing the conformational entropy of unfolding (23, 37, 40), which is known as the “proline theory.” From the sequence homology among the four chitinases (Figure 1), we noticed the five residues where proline residue is conserved were in some other chitinases but not in GlxChiB, and we designed five mutants (A117P, A151P, Q171P, A233P, and A254P). All five mutants and integrated mutant, mt5, exhibited a positive effect on the thermostability (Table 2 and Figure 6). From the structural analysis (Figure 4), it was clarified that Ala117 is located at the second site of beta-turn I, Ala151 is located at the second site of beta-turn II, Gln171 is in the conserved helix structure, Ala233 and Ala254 are located at the loop region preceding conserved helix structure in CD of WT-GlxChiB, and Ala254 is located at N-terminal caps of α-helices. Proline residues contributing to thermal stabilization favor second sites of β-turns, loop regions, and N-terminal caps of α-helices (23, 36). Therefore, the construction of the thermostable mutants (A117P, A151P, A233P, and A254P) by the substitution of proline is well explained as the proline theory (36, 41).

Q171P is in a different circumstance; the residue is located in the middle of the alpha-helices structure, and proline residue is well known as the alpha-helices breaker (28). However, the Gly170 preceding the Pro171 appears to release the distorted structure by Pro171 (Figure 4C). In addition, Tyr147 and Trp200 exhibit the hydrophobic interaction toward Pro171 (Figure 7). The hydrophobic interaction appears to contribute to the thermostability of Q171P. For position 147, aromatic residues (Tyr, Phe, and Trp) are well conserved in the other chitinase, and Trp200 is also conserved in the enzyme (Figure 1). It is noteworthy that this hydrophobic interaction strengthens the connection between two helices and a loop region, contributing to the thermostability.

**Figure 7.**
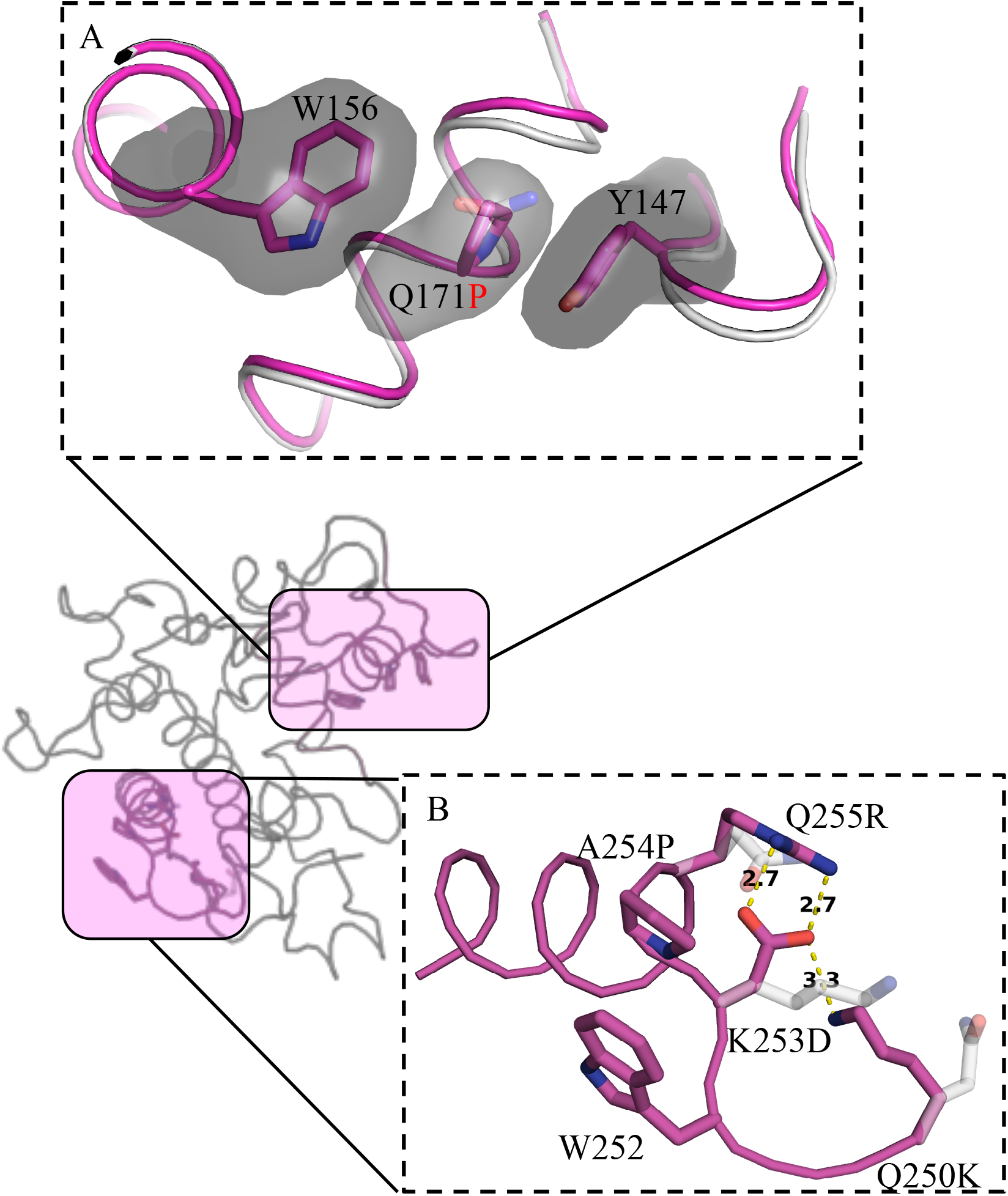
New interactions in thermostable mutants and the estimated salt bridges. A, The hydrophobic interactions toward Pro171. The surface of Pro residue and two aromatic residues, Y147 and W156 were shown in gray. B, Structure of loop region from Q250 to Q255 in mt5. The salt bridge network among K250, D253, and R255 was estimated using a PyMOL mutagenesis option based on the structure of mt5 (the orientation of the side chains is adjusted to avoid steric hindrance). Side chains of original residues, Q250, K253, and Q255 are shown as stick model colored transparent. The Estimated hydrogen bonding among K250, D253, and R255 are indicated as yellow dashed lines with labeled number as the distance (Å).

### Structural interpretation of the thermostability by SS bond

A disulfide bridge linking the two cysteines stabilizes the folded state of the protein. From the crystal structural data of CD-WT and the sequence homology among the enzymes, we cannot find the suitable position in which the additional disulfide bonding can be introduced in the catalytic domain. However, it was estimated that the disulfide bonding between Gly44 (in the linker region) and Lys83 could form. In the DSC analysis, the melting temperature (Tm) of the mutant G44C/K83C was not higher than that of WT. However, the half-life at 60 °C was elongated without depressing their activities (Table 2 and Figure 6). By introducing the disulfide bonding, the thermostability of the enzyme is not improved; however, irreversible inactivation appears to be suppressed.

### Structural interpretation of the thermostability by Salt bridge

It is well known that salt bridges contribute to the stability of proteins (42, 43). Ideally, the introduction of salt bridges can increase the thermostability of the enzyme. However, the fundamental problem is that the formation of salt bridges depends on the ionization properties of the participating groups, which are significantly influenced by the environmental changes around the proteins (44). Furthermore, salt bridges experience thermal fluctuations, continuously break and reform, and the flexibility of the protein governs their lifespan in solution. Nevertheless, proteins from thermophiles and hyperthermophiles exhibit more frequently networked salt bridges than proteins from the mesophilic counterparts (43, 45, 46). Increasing the thermostability of proteins by optimizing charge-charge interactions is a good example of an evolutionary solution utilizing physical factors. For the two mutants with a salt bridge network introduced (Q250K/K253D/Q255R and Q250R/K253D/Q255R), specific activities at 37 °C were slightly increased. In addition, the half-life time at 60 °C was increased (Table2 and Figure 6). Figure 7B shows a model structure of the loop region (250Q-255Q) between two helices in the catalytic domain of mt5ss/Q250K/K253D/Q255R based on the crystal structure of mt5. It is speculated that the salt bridge networks among Lys250, Asp253, and Arg255 are constructed by inferring the salt bridge network in other GH19 structures (Figure S2). It should be noted that this region involves the proline substitution site (A254P). Thus, the performance of mt5ss/Q250K/K253D/Q255R is achieved by multiple effects of the hydrophobic interaction between Trp252 and Pro254, the stabilized peptide chain by Pro254, and the salt bridge network among Lys250, Asp253, and Arg255. In addition, Asp253 in mt5ss/Q250K/K253D/Q255R appears to form ASX turn preceding α-helices (Figure 1 and 7B) (47).The α-helix has an overall dipole moment due to the aggregate effect of the individual microdipoles. Therefore, α-helices often occur with the N-terminal end bound by a negatively charged group (48). It is speculated that Asp253 (negative charge) in mt5ss/Q250K/K253D/Q255R also increases the stability of the α-helices in the enzyme. We cannot explain why the thermostability of mt5ss/Q250H/K253D/Q255R is better than mt5ss/Q250K/K253D/Q255R. However, this 250^th^ histidine residue is conserved in some chitinases (Figure 1), and an additional mechanism of the thermostability seems to be involved. The detailed study for the position is in progress.

### Thermostable mutation effects on the catalytic and antifungal activity

The trade-off between the stability and function of the enzyme is widely recognized from the observation that the substitution of catalytic residues can dramatically improve its stability at the expense of its activity (29-31). Improvement of thermostability without reducing enzyme activity is highly demanded in industrial applications. We targeted the mutation sites that are apart from the catalytic residues (Figure 3). As a result, all mutations applied to GlxChiB in this study did not reduce its catalytic activity and antifungal activity (Table 2). Furthermore, some mutants exhibit higher antifungal activity than WT. It has been reported that decreasing the conformational entropy of unfolding contributes to a resistance to protease (49-51). Therefore, some thermostable mutations appear to affect the activity itself but improve the resistance to protease secreted from the fungus, resulting in the improvement of the antifungal activity. In summary, we have succeeded in making the enzyme thermostable without decreasing its high catalytic activity and antifungal activity by using protein engineering methods. This study provides a successful strategy to improve the thermostability of GH19 chitinase and identifies the thermostable mutants of GlxChiB as a good seed for industrial applications.

## MATERIALS AND METHODS

### Construction of mutant protein genes

The genes encoding GlxChiB and its mutants were cloned into pET22b (Novagen) at the NdeI and BamHI restriction sites. All mutant genes were constructed by polymerase chain reactions using a QuickChange Site-Directed Mutagenesis kit (Stratagene) and primers (Table S1). Confirmation of the plasmid DNA sequences was outsourced to Fasmac Japan (Kanagawa, Japan).

### Expression and purification of the recombinant protein

Plasmids encoding GlxChiB and its mutants were transformed into *E*.*coli* SHuffle T7 (DE3) cells (New England Biolabs). Cells harboring the plasmids were cultured at 37 °C in LB medium containing 100 mg/L ampicillin-Na for about 3.5 hours to reach OD600 of 0.6-0.8 and then were induced to express recombinant protein by adding IPTG to a final concentration of 0.1 mM. The culture was incubated for an additional 60 hours at 18 °C, and then the cells were harvested and disrupted by sonication in 20 mM Tris-HCl buffer, pH 8.0. The sonicated extract was separated into soluble and insoluble fractions by centrifugation at 12,000×g for 15 minutes at 4 °C. The soluble fraction was dialyzed against 10 mM sodium phosphate buffer, pH 7.0, and apply to a RESOURCE S column (6 mL, GE Healthcare) previously equilibrated with the same buffer. The elution was done with a linear gradient of NaCl from 0 to 0.3 M in the same buffer. The fractions containing recombinant protein were collected and dialyzed against 10 mM sodium phosphate buffer, pH 7.0. The purity of the recombinant protein was analyzed by SDS-PAGE by the Laemmli method (52) using 12.5% polyacrylamide gels.

### Protein assay

All protein concentrations were determined with the bicinchoninic acid (BCA) method (53). The protein concentration was determined with the Pierce BCA Protein Assay Kit (Thermo Scientific) using bovine serum albumin as the protein standard.

### Chitinase activity assay

Chitinase activity was assayed colorimetrically with glycol chitin as a substrate. Glycol chitin was prepared by the method described by Yamada & Imoto (54). Ten microliters of the sample solution were added to 250 μL of 0.2% (w/v) glycol chitin solution in 0.1 M sodium phosphate buffer, pH 7.0. After incubation of the reaction mixture at 37 °C for 15 minutes, the reducing power of the mixture was measured with ferric ferrocyanide reagent by the method of Imoto & Yagishita (55). One unit of activity was defined as the enzyme activity that produced 1 μmol of GlcNAc per minute at 37 °C. The thermal stabilities of the enzymes were assessed by measuring the residual activities after incubation in 10 mM sodium phosphate buffer, pH 7.0 at 60 °C and 65 °C for the appropriate length of time. The residual activities were measured under the standard condition.

### Quantitative antifungal activity assay (Taira et al., 2002) (56)

Hyphal re-extension inhibition assay was done by using *Trichoderma viride*. Agar disks (4 mm in diameter) containing the fungal hyphae, which were derived from the resting part of the fungus previously cultured on potato dextrose broth containing 1.5% (w/v) agar (PDA), were put on another PDA plate with the hyphae attached side down. Five microliters of sterile water or sample solution were overlaid on the agar disks, and then the plate was incubated at 25 °C. for 12 hours. After incubation, images of the plates were scanned using an image scanner. The areas of the re-extended hyphae were calculated as numbers of pixels by GNU Image Manipulation Program (GIMP, ver. 2.0). The protein concentration required for inhibiting the growth of the fungus by 50% was defined as IC_50_ and determined by constructing dose-response curves (percentage of growth inhibition versus protein concentration).

### Differential scanning calorimetry (DSC)

The thermal stability of the enzymes was examined using differential scanning calorimetry (DSC). GlxChiB or its mutants in 10 mM sodium phosphate buffer (pH 7.0) were used at a final concentration of 1.0 mg/mL. A Nano DSC instrument (TA Instruments) was used at a scanning speed of 60 °C/h. Control runs in the absence of protein were carried out before and after each sample run. DSC scans in the presence of protein were performed two or three times for each protein examined.

### Crystallization

The recombinant catalytic domain of the enzyme was prepared and purified as described previously (7). The purified proteins were dialyzed against 5 mM Tris-HCl buffer (pH 8.0) and concentrated to 10 mg/mL. Initial crystallization screening of the mutant protein was performed using various crystallization screening kits commercially available. The protein solution drop (0.15 μL) was mixed with 0.15 μL of a reservoir solution and then equilibrated with 50 μL of the reservoir solution. The crystallization was carried out according to the hanging-drop vapor diffusion method at 293 K in 96-well plates. After a week, well-formed crystals of CD-WT were obtained from the optimized condition (3% (w/v) gamma-polyglutamic acid low molecule, 25%(w/v) 2-Methy-2, 4-pentanediol, 0.1 M HEPES (pH 7.5), and 0.5 M Ammonium sulfate). Well-formed crystals of CD-mt5 were obtained from the different condition than CD-WT (18% (w/v) Polyethylene glycol 20K, 0.1 M sodium citrate (pH 5.0), and 3% (v/v) glycerol).

### X-ray data collection

The crystal of CD-WT and CD-mt5 were frozen in liquid nitrogen. The diffraction data sets were collected at a wavelength of 1.000 Å at BL5A and NE3A beamline of the Photon Factory in KEK, Japan, respectively. Data was processed by the program HKL2000 (57) for CD-WT and XDS (58) for CD-mt5.

### Structure solution and refinement

General data handling was carried out with the CCP4 package (59). The initial model was solved by molecular replacement using Phaser (60) with the crystal structure of chitinase (GH19) from *Vigna unguiculata* (PDB: 4TX7) (17) as the search model for CD-WT. The model building was carried out with *Coot* (61) and refinement using REFMAC5 (62). Structural figures are described and rendered by the PyMOL Molecular Graphics System, Version 1.2r3pre, Schrödinger, LLC.

## Acknowledgments

This work was performed as a part of Projects of Okinawa innovation system building business of science and technology, supported by the Okinawa Science and Technology Promotion Center.

## Author contributions

DK, HF and TK performed the experiments. DK and KI wrote the manuscript. KI, TT, TK, and KU thoroughly revised the manuscript. KI and TT designed and supervised the project. All authors read and approved the manuscript.

## Supplemental information

**Table S1.**
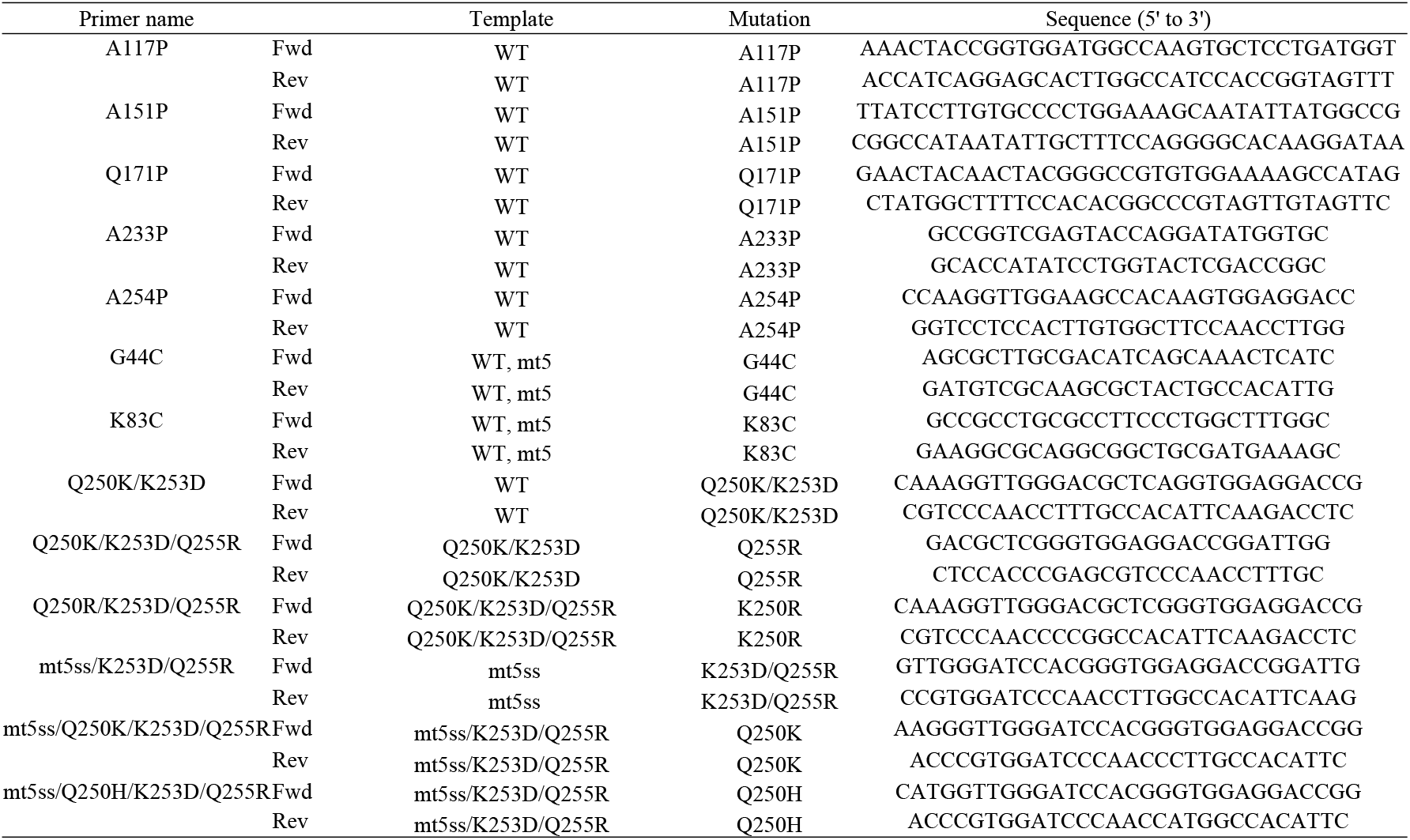
Site-directed mutagenesis primers.

**Figure S1.**
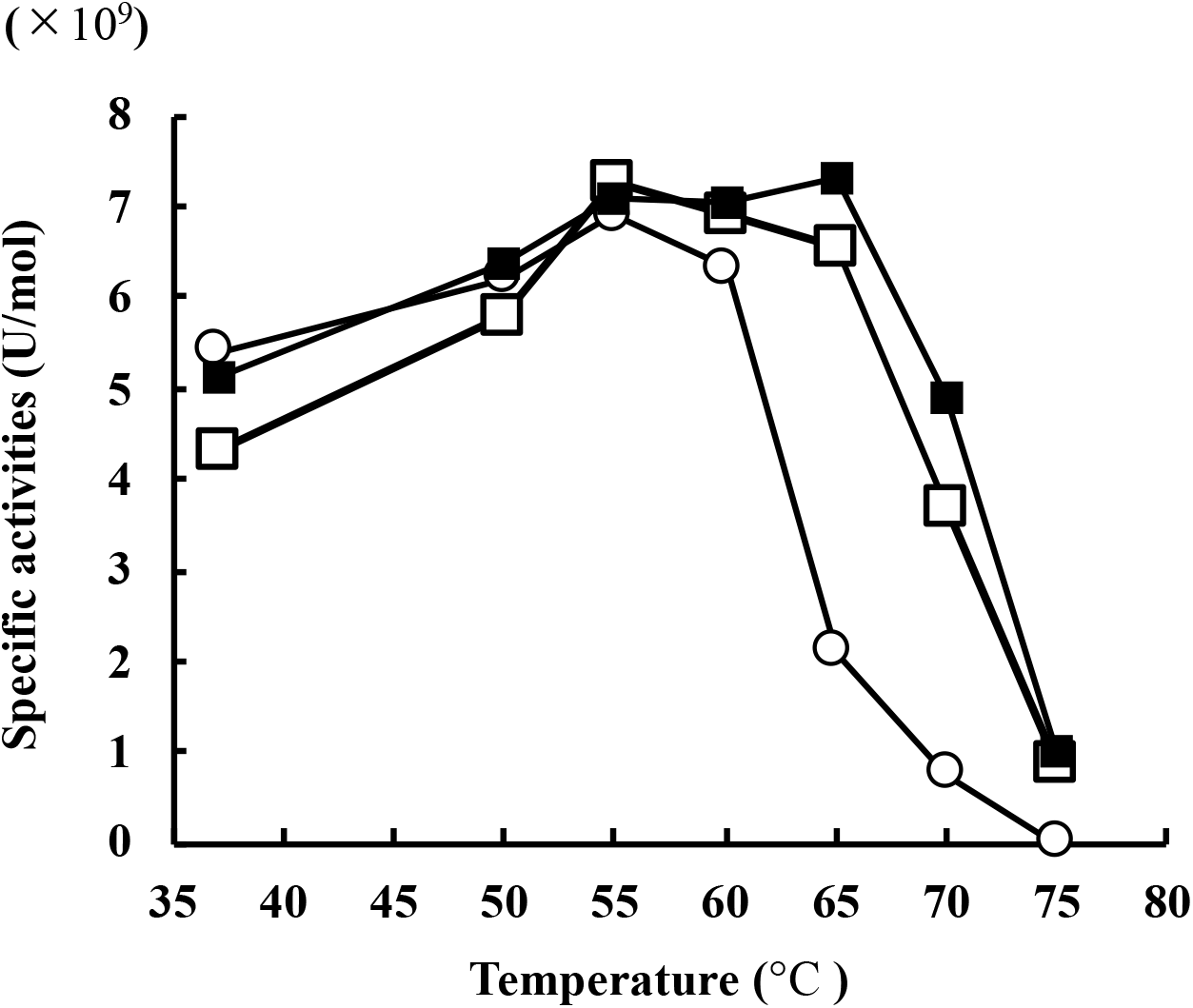
Optimum temperature of WT and integrated mutants. Specific activities were determined using 0.2% (w/v) glycol chitin as substrate in 0.2 M sodium phosphate buffer (pH 7.0) for 15 minutes. Open circles indicate WT. Open and closed squares indicate mt5ssKDR and mt5ssHDR, respectively.

**Figure S2.**
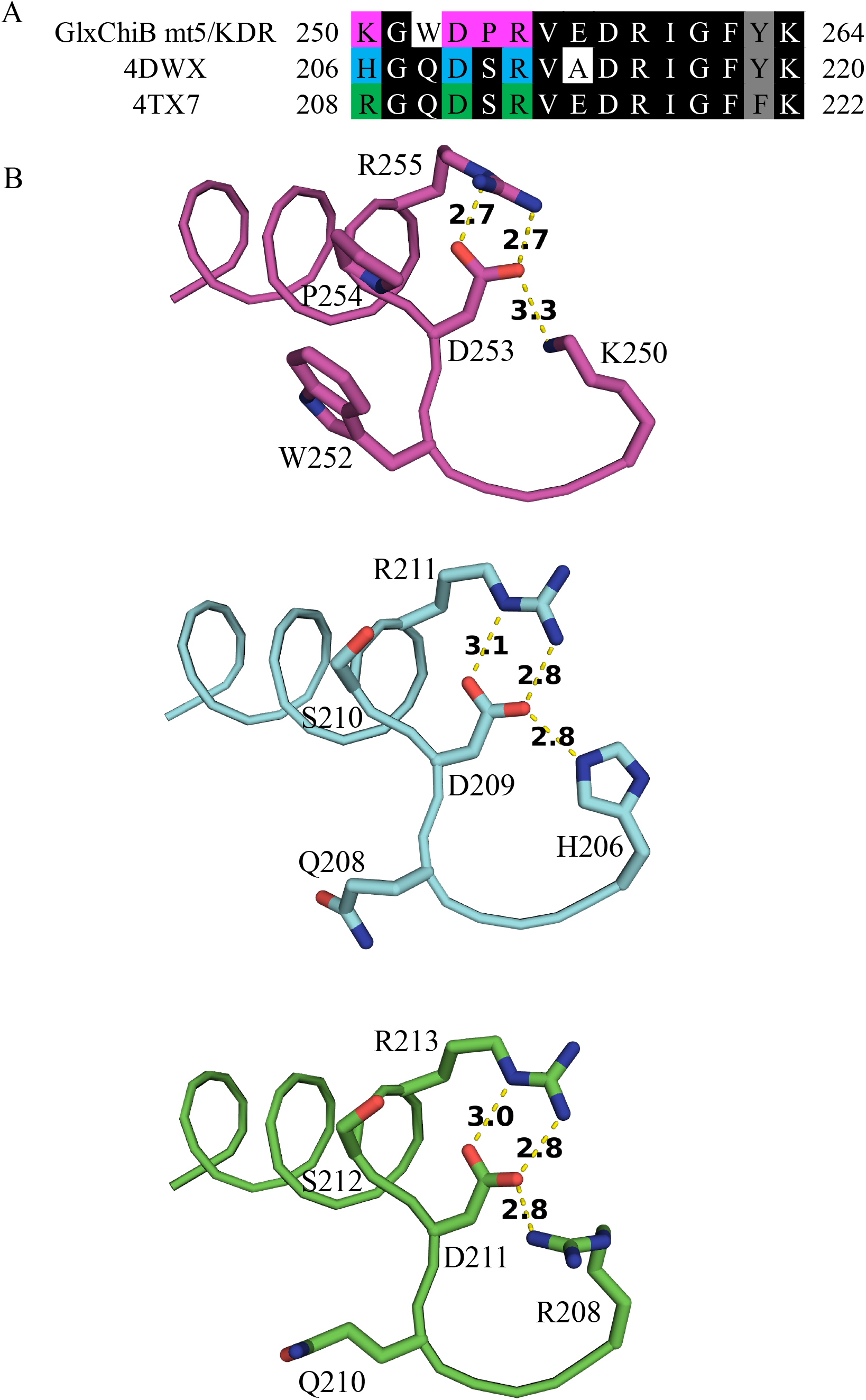
comparison of salt bridge network in other GH19 chitinases. A, Multiple sequence alignment among GlxChiB mt5/KDR, 4DWX, and 4TX7. B, Structural comparison of the salt bridge network among these three GH19 chitinases.

